# Self-reversal facilitates the resolution of HMCES-DNA protein crosslinks in cells

**DOI:** 10.1101/2023.06.14.544844

**Authors:** Jorge Rua-Fernandez, Courtney A. Lovejoy, Kavi P.M. Mehta, Katherine A. Paulin, Yasmine T. Toudji, Brandt F. Eichman, David Cortez

## Abstract

Abasic sites are common DNA lesions that stall polymerases and threaten genome stability. When located in single-stranded DNA (ssDNA), they are shielded from aberrant processing by HMCES via a DNA-protein crosslink (DPC) that prevents double-strand breaks. Nevertheless, the HMCES-DPC must be removed to complete DNA repair. Here, we found that DNA polymerase α inhibition generates ssDNA abasic sites and HMCES-DPCs. These DPCs are resolved with a half-life of approximately 1.5 hours. Resolution does not require the proteasome or SPRTN protease. Instead, HMCES-DPC self-reversal is important for resolution. Biochemically, self-reversal is favored when the ssDNA is converted to duplex DNA. When the self-reversal mechanism is inactivated, HMCES-DPC removal is delayed, cell proliferation is slowed, and cells become hypersensitive to DNA damage agents that increase AP site formation. Thus, HMCES-DPC formation followed by self-reversal is an important mechanism for ssDNA AP site management.

## INTRODUCTION

Abasic sites are one of the most common DNA lesions. They can be formed by spontaneous depurination/depyrimidination or as intermediates during the excision of damaged nucleobases by glycosylases (Nakamura and Nakamura 2020; Thompson and Cortez 2020; Friedberg et al. 2006). When AP sites are within double-stranded DNA (dsDNA), they can be repaired by base excision repair (BER). The endonuclease APE1 cleaves 5’ to the AP site and Polβ uses the intact DNA strand as a template for synthesis to complete repair (Krokan and Bjoras 2013). AP sites can also be converted into strand-breaks via a spontaneous β-elimination reaction (Thompson and Cortez 2020; Nakamura and Nakamura 2020). Spontaneous or enzymatic cleavage of AP sites located in ssDNA can lead to double-strand breaks (DSBs), which are highly toxic to cells. During DNA replication, AP sites are potent blocks to replicative polymerases placing them at a dsDNA-ssDNA junction (Choi et al. 2010). Translesion DNA synthesis (TLS) polymerases can synthesize DNA across AP sites but often will cause mutations.

HMCES (5-hydroxymethyl cytosine, embryonic ES-cell-specific) was recently identified as a shield for ssDNA AP sites (Mohni et al. 2019). HMCES is present at replication forks, interacts with PCNA, and covalently binds to ssDNA AP sites through an evolutionary conserved SOS-response associated peptidase (SRAP) domain, generating a DPC (Mohni et al. 2019; Srivastava et al. 2020). Unlike damaging DPC formation with other proteins (Nakamura and Nakamura 2020), the HMCES-DPC is thought to be protective and beneficial to the cell. The HMCES-DPC prevents AP site cleavage thereby reducing DSBs (Mohni et al. 2019; Thompson et al. 2019) and also decreases mutation frequency (Srivastava et al. 2020; Mohni et al. 2019). HMCES and its bacterial ortholog YedK form a thiazolidine linkage between a ring-opened AP site and an N-terminal SRAP cysteine residue (Cys 2) (Thompson et al. 2019; Wang et al. 2019; Halabelian et al. 2019). Inactivating HMCES through mutation of the cysteine, RNA interference, or gene disruption causes hypersensitivity to DNA damaging agents that increase AP site frequency (Mohni et al. 2019; Srivastava et al. 2020) and synthetic lethality with nuclear expression of cytidine deaminase APOBEC3A (Biayna et al. 2021; Mehta et al. 2020), which also increases AP site formation (Mehta et al. 2020). HMCES is also important to suppress deletions during somatic hypermutation (SHM) (Wu et al. 2022) and class switch recombination (Shukla et al. 2020) in B cells. Thus, HMCES acts by shielding ssDNA AP sites from inappropriate processing. Nevertheless, the HMCES-DPC itself is a bulky lesion that can interfere with replication and transcription (Sugimoto et al. 2023). Persistence of DPCs in cells is associated with different diseases (Ruggiano and Ramadan 2021). Thus, the protective activity of HMCES-DPC should include its removal.

The HMCES-DPC is ubiquitylated, and proteasome inhibition may delay DPC removal after potassium bromate (KBrO_3_) treatment (Mohni et al. 2019), suggesting a proteasome-dependent degradation of the DPC. In the *Xenopus* egg extract system, the HMCES-DPC is removed by the SPRTN protease as an intermediate step in a DNA interstrand crosslink (AP-ICL) repair pathway that generates a ssDNA AP site (Semlow, MacKrell, and Walter 2022). Proteasome inhibition had no effect in this system, despite HMCES-DPC being a target for ubiquitylation by the E3 ligase RFWD3 in *Xenopus* egg extracts (Gallina et al. 2020). Other proteases have been reported to remove DPCs to maintain DNA replication and genome stability (Borgermann et al. 2019; Kojima et al. 2020; Yip, Bodnar, and Rapoport 2020); however, whether any act on HMCES-DPCs is unknown.

Recently, the HMCES-DPC and YedK-DPC were shown to be reversible in biochemical reactions (Paulin, Cortez, and Eichman 2022; Sugimoto et al. 2023; Donsbach et al. 2022). Two residues in close proximity to the thiazolidine linkage, His160 and Glu105, in YedK are important for this crosslink reversal process (Paulin, Cortez, and Eichman 2022). Mutation of the human HMCES equivalent of Glu105 (Glu127) also impairs HMCES-DPC reversal *in vitro* (Donsbach et al. 2022). It is unknown whether this self-reversal activity is biologically important. Here, we developed a cellular system to detect, quantify, and track HMCES-DPC resolution. Our results provide evidence that self-reversal is an important mechanism in cells to remove the HMCES-DPC and promote cell fitness.

## RESULTS

### POLα inhibition generates ssDNA AP sites and HMCES-DPCs

Previous cellular studies of HMCES-DPC formation utilized DNA damaging agents like KBrO_3_ to generate abasic sites. These approaches yield a modest increase in HMCES-DPC levels (approximately 3-fold) (Mohni et al. 2019). The generation of abasic sites in these circumstances occurs mostly in duplex DNA. Formation of the HMCES-DPC in response to this DNA damage likely requires unwinding of the DNA during DNA replication to move the AP site into ssDNA where HMCES functions. Therefore, both the generation and resolution of the HMCES-DPC in response to these types of DNA-damaging agents take hours. To better study DPC resolution we looked for a cellular system in which AP site formation would be targeted to ssDNA and can be better separated from DPC resolution. CD437 is a direct polymerase alpha (POLα) inhibitor (Han et al. 2016) that prevents the synthesis of the lagging strand and rapidly generates large amounts of ssDNA compared to other replication stalling agents like hydroxyurea (Figure 1A) (Ercilla et al. 2020). ssDNA is more vulnerable to chemical attack and spontaneous depurination (Chan et al. 2012), which leads to AP site formation. Indeed, we could detect AP sites with an aldehyde reactive probe (ARP) after the addition of CD437 (Figure 1B). Since HMCES crosslinks specifically to AP sites located in ssDNA (Mohni et al. 2019), we hypothesized that CD437 would induce HMCES-DPC formation. As expected, cells treated with 5 μM CD437 for 30 min showed a 10-fold increase in HMCES-DPC signal using the rapid approach to DNA adduct recovery (RADAR) assay (Figure 1C) (Kiianitsa and Maizels 2013). Moreover, this signal was reduced in cells transduced with a plasmid expressing the uracil DNA glycosylase inhibitor (UGI) (Cone, Bonura, and Friedberg 1980) (Figure 1C), confirming HMCES reacted with AP sites largely created by glycosylase activity in these conditions.

**Figure 1.**
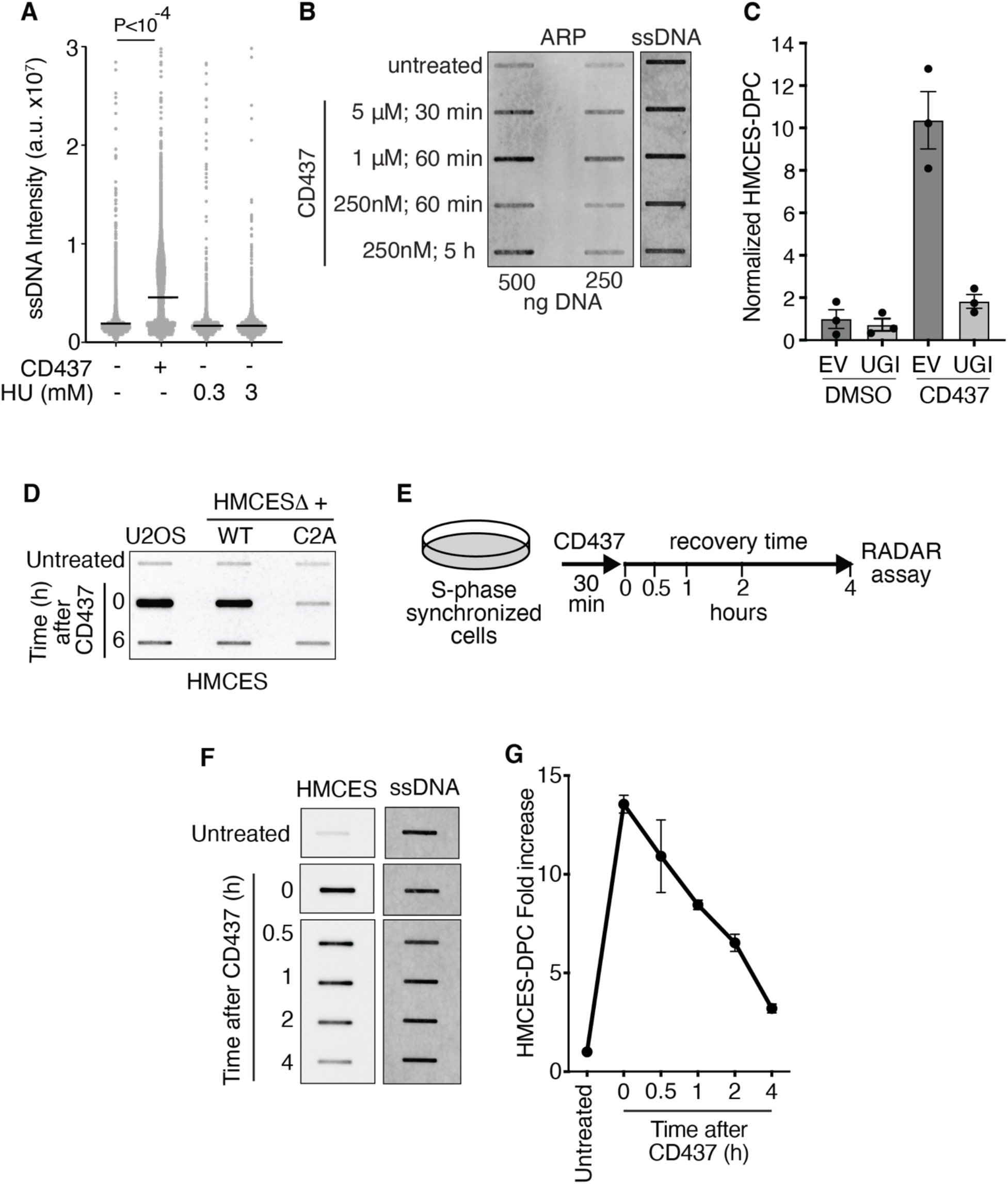
CD437 induces AP sites in ssDNA and HMCES-DPC formation in cells. **(A)** Levels of ssDNA in HCT116 cells after treatment with 5 μM CD437, 0.3mM or 3mM HU for 30 min. Intensity from individual nuclei and mean is shown, Kruskal-Wallis test. **(B)** Aldehyde reactive probe (ARP) assay on HCT116 cells that were exposed to drugs as indicated. **(C)** Quantification of HMCES-DPC using RADAR assay. Cell expressing empty vector (EV) or UGI were treated with 5 μM CD437 or DMSO for 30 min. **(D)** Representative HMCES-DPC assay in U2OS or HMCESΔ cells complemented with WT or C2A HMCES. **(E)** Schematic for detection of HMCES-DPC removal in cells after CD437 treatment. **(F)** Representative image of HMCES-DPC resolution assay. Cells were synchronized and released from thymidine for 2 h prior to treatment with CD437 for 30 minutes. Samples were taken immediately after CD437 treatment (0) and at the indicated time points. **(G)** Quantification of HMCES-DPC resolution. Mean ± SEM, n=3.

To confirm that CD437-induced HMCES-DPCs are formed via a thiazolidine linkage, we tested DPC formation in cells in which endogenous *HMCES* was deleted by gene editing (HMCESΔ) and either wild-type (WT) or a HMCES C2A mutant protein that is unable to crosslink to the abasic site was expressed by retroviral integration. As expected, no HMCES-DPCs were detected in cells expressing HMCES C2A after CD437 treatment; whereas cells complemented with wild-type HMCES had an equivalent DPC level as the parental U2OS cells (Figure 1D). The HMCES C2A protein was expressed at similar levels to WT (Figure S1).

We next asked whether the HMCES-DPCs induced by CD437 were resolved over time. S-phase synchronized cells were treated with CD437 for 30 minutes to induce HMCES-DPC formation. After removing CD437, cells were incubated in normal growth media and harvested at varying time points to analyze HMCES-DPC levels (Figure 1E). We observed a strong HMCES-DPC signal immediately after CD437 treatment that declined rapidly during recovery (Figure 1F). Quantification shows a HMCES-DPC half-life between one and two hours and approximately 80% of the HMCES-DPC is removed after 4 hours (Figure 1G). We conclude that CD437 promotes HMCES-DPC formation in cells by increasing AP sites in ssDNA, and provides a quantifiable system to analyze HMCES-DPC removal.

### CD437-induced HMCES-DPC removal does not require the proteasome or SPRTN protease

Proteolysis is a major pathway to repair DPCs during DNA replication (Kühbacher and Duxin 2020), and treatment with a proteasome inhibitor (MG132) appeared to delay HMCES-DPC resolution after KBrO_3_ treatment (Mohni et al. 2019) suggesting a proteasome-dependent degradation of the HMCES-DPC. To test the activity of the proteasome in HMCES-DPC removal, we utilized our CD437 system to track DPC levels in the absence or presence of MG132. Treating HCT116 cells with MG132 increased the total amount of HMCES-DPC formed after CD437 incubation by approximately 25% 30 minutes after CD437 removal (Figure 2A). However, MG132 did not prevent the resolution of HMCES-DPCs which proceeded at least as quickly after the 30-minute time point as vehicle-treated cells (Figures 2A and S2A). Incubating cells with MG132 alone in the absence of CD437 does not induce HMCES-DPC formation (Figure S2B); therefore, the increase within the first 30 min of release from CD437 required ssDNA formation. We also observed similar results by inhibiting the UBA1 E1 enzyme with TAK243 (Barghout et al. 2019) which blocks protein ubiquitylation (Figures 2B, S2A, and S2B). These results suggest that ubiquitylation and the proteasome are not absolutely essential for resolving the HMCES-DPC, although they may affect the amount of DPC formed or retained at early time points after CD437 treatment.

**Figure 2.**
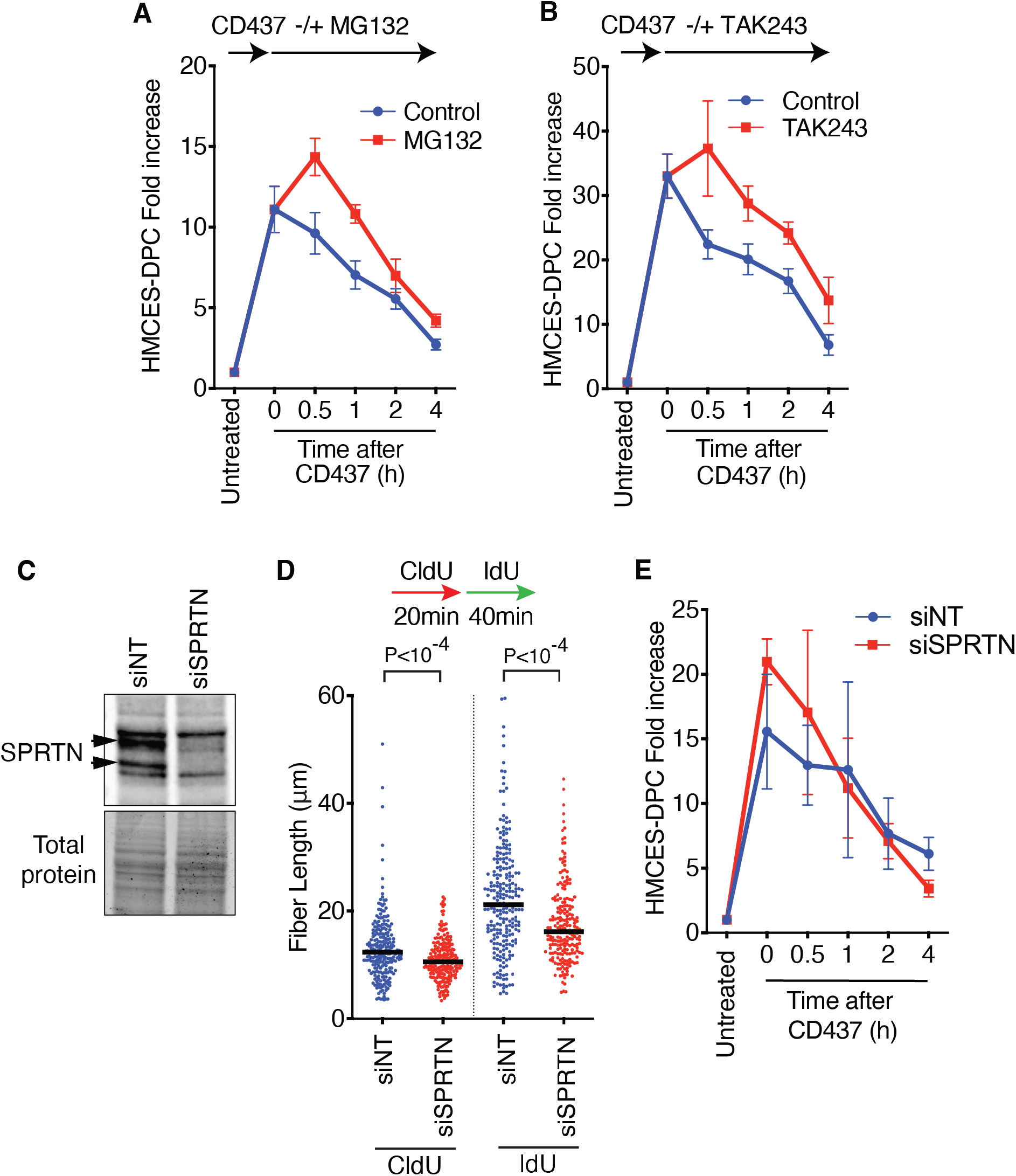
Inactivation of the proteasome or SPRTN does not prevent HMCES-DPC removal. **(A)** Quantification of HMCES-DPC removal. MG132 was added after CD437 treatment. Mean ± SEM, n=5. **(B)** Quantification of HMCES-DPC removal. TAK243 was added after CD437 treatment. Mean ± SEM, n=4. **(C)** Immunoblot using SPRTN antibody of cells transfected with no-targeting siRNA (siNT) or siRNA against SPRTN (siSPRTN). **(D)** DNA combing assay analysis of replication fork speed in cells transfected with non-targeting or SPRTN siRNA. Bar represents the median. P values derived from Mann-Whitney test. **(E)** Quantification of HMCES-DPC removal in cells transfected with siRNA, Mean ± SEM, n=3.

We next tested whether the SPRTN protease is important for HMCES-DPC resolution. SPRTN is essential for mammalian cell survival (Maskey et al. 2014); therefore, we used siRNA to deplete SPRTN acutely and test its activity. Knockdown efficiency was confirmed by immunoblotting (Figure 2C). We also verified that SPRTN was functionally inactivated by measuring replication fork speed, which was previously shown to be slowed by SPRTN inactivation (Figure 2D) (Halder et al. 2019). Depletion of SPRTN did not prevent CD437-induced HMCES-DPC removal (Figure 2E). Additionally, we tested a possible redundancy between SPRTN and the proteasome; however, MG132 did not prevent HMCES-DPC removal in SPRTN knockdown cells (Figure S2C). Therefore, our data suggest that neither the proteasome nor SPRTN activity is critical to remove the HMCES-DPC induced by POLα inhibition.

### HMCES Glu127 mediates the self-reversal reaction which is stimulated by a duplex forming oligonucleotide

Although the HMCES-DPC thiazolidine linkage appears to be stable and resistant to repair enzymes like AP endonucleases, biochemical experiments using a second ssDNA containing an AP site as a trap showed that it is reversible (Paulin, Cortez, and Eichman 2022). In the HMCES bacterial ortholog YedK, the reversal reaction was disrupted in E105Q and H160Q mutants with the former having a stronger effect. To test if the equivalent glutamic acid residue is also important for the self-reversal of human HMCES, we took a similar experimental approach. The HMCES-DPC was formed by incubation of the SRAP domain with a 20 nucleotide DNA oligo containing an AP site (DPC-20). Next, we added a 40 nucleotide AP DNA oligo in 50-fold excess to trap any self-reversed protein during incubation at 37 degrees (Figure 3A). The DPC-20 slowly decreased over time as a DPC-40 formed in accordance with self-reversal regenerating an intact and active HMCES protein capable of crosslinking again to another available ssDNA AP site (Figure 3B). In contrast, the E127Q HMCES-DPC is completely unable to reverse even after 24 hours of incubation. This result indicates that E127 is necessary for human HMCES-SRAP crosslink self-reversal.

**Figure 3.**
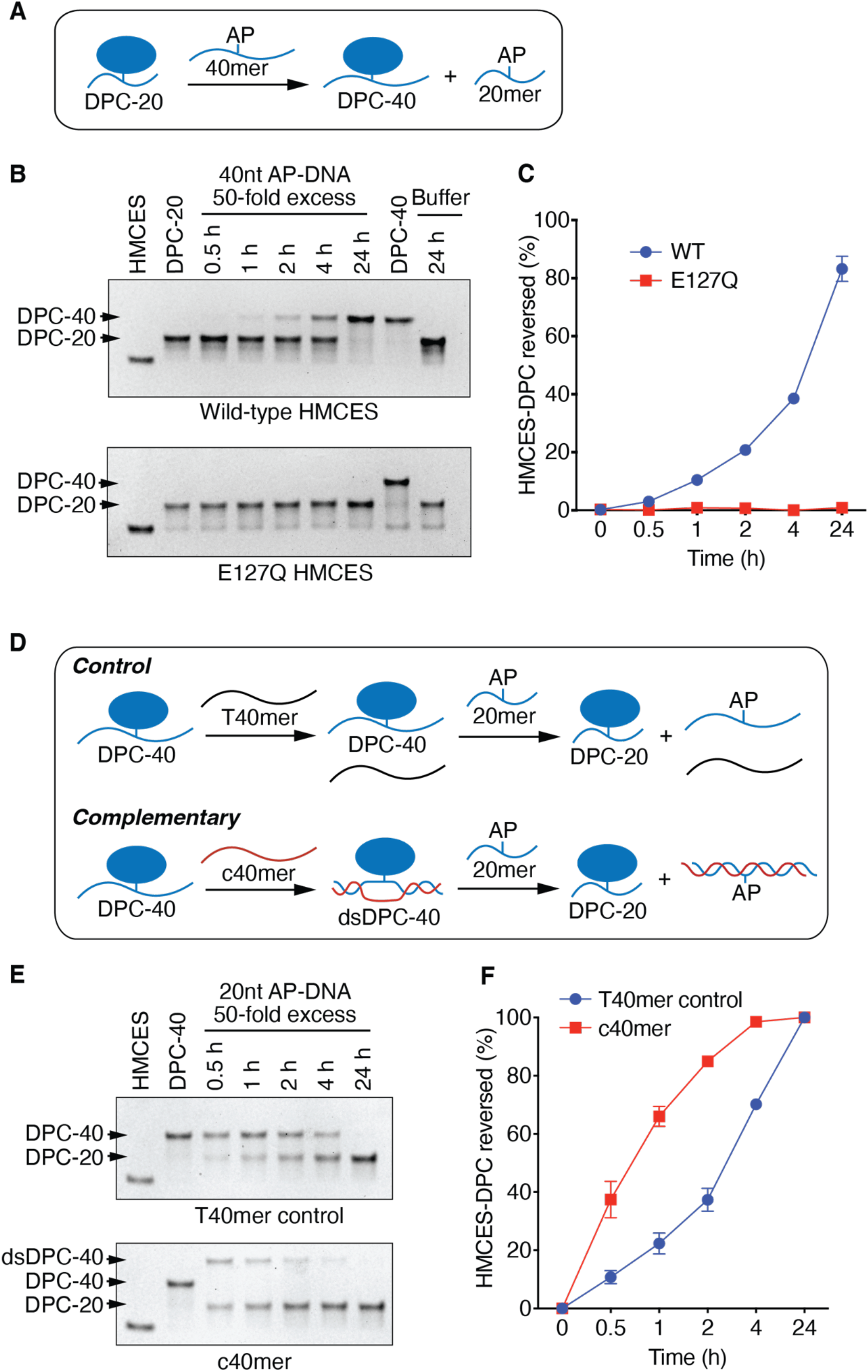
HMCES-DPC self-reversal is dependent on E127 and is stimulated by duplex DNA. **(A)** Schematic of HMCES-DPC biochemical self-reversal assay. The AP site containing 40 nucleotide ssDNA oligonucleotide (40mer) was added at 50x excess. **(B)** Representative time course experiment of HMCES-DPC reversal with purified wild-type or E127Q HMCES. The first lane is HMCES protein without any DNA. The free protein and DPC-20 (DPC with 20 mer) or DPC-40 (DPC with 40 mer) were visualized by coomassie-stained SDS-PAGE. **(C)** Quantification of HMCES-DPC reversal. Percent reversed is the amount of DPC-40 product compared to the total. Mean ± SEM, n=3. **(D)** Schematic of HMCES-DPC reversal assay comparing single-stranded vs duplex DNA. The HMCES DPC-40 was incubated with a complementary (c40mer) or non-complementary (T40mer control) oligonucleotide with a 20 nucleotide ssDNA trap containing an AP site. **(E)** Representative time course of HMCES-DPC reversal. The higher migrating band is the product of the hybridization of c40mer with ssDNA HMCES-DPC (dsDPC-40). **(F)** Quantification of HMCES-DPC reversal from ssDNA and dsDNA. Percent reversed is the amount of the DPC-20 product compared to the total. Mean ± SEM, n=3.

The half-life of the HMCES-DPC self-reversal *in vitro* is longer than four hours (Figure 3C). This is much greater than the half-life of the DPC seen after CD437 treatment in cells. Moving from one AP-ssDNA site to another requires not only the reversal of the thiazolidine linkage but also the disengagement of DNA binding. Our previous studies showed that even without crosslinking, HMCES has a very high affinity for ssDNA (Mohni et al. 2019). In contrast, HMCES cannot bind dsDNA. Thus, we reasoned that if a complementary oligonucleotide capable of forming a duplex with the ssDNA to which HMCES is crosslinked was included in the reversal reaction, we may be able to increase the speed at which we could observe the reversal. Indeed, this is the case. After generating HMCES-DPC-40, we added a complemented oligo (c40mer), which shifted the DPC-40 band in an SDS-PAGE gel (dsDPC-40). Next, we added a 50-fold excess of 20mer containing an AP site to trap any HMCES that undergoes self-reversal (Figure 3D). As expected, the control sample containing a non-duplex forming 40mer oligonucleotide (T40mer control) showed a reduction of DPC-40 along with the formation of DPC-20. However, the reversal of DPC-40 in the presence of the duplex-forming oligonucleotide is faster compared to the control (Figure 3E). After one hour incubation, the dsDNA structure yielded more than 60% reversal, compared to HMCES-DPC in ssDNA, which reached a little more than 20% (Figure 3F). This result is consistent with two recent reports that found that DNA duplex formation caused an apparent accelerated self-reversal rate (Sugimoto et al. 2023; Donsbach et al. 2022) and indicates that the intrinsic rate of self-reversal is comparable with the reversal rate observed in cells.

### HMCES-DPC self-reversal is an important resolution pathway in human cells

To test if HMCES-DPC auto-reversal is an important resolution pathway in cells, we made use of the E127Q HMCES protein. First, we complemented the HMCESΔ cells to create stable cell lines expressing only the E127Q or WT HMCES (Figure 4A). Analysis of HMCES-DPC levels in untreated, synchronized cells showed more DPC in E127Q cells compared to WT cells (Figure 4B), suggesting that the crosslinked state is increased in cells expressing the mutant even in the absence of added genotoxic stress. We next treated the E127Q or WT HMCES expressing cells with CD437 to induce HMCES-DPC formation and tracked DPC resolution over time. The E127Q HMCES-DPC formed a similar total level of DPC as wild-type cells 30 minutes after CD437 treatment (Figure 4C), although the fold-increase when compared to the untreated cell control was less since it started at a higher basal level. Strikingly, the resolution kinetics of the E127Q was significantly delayed by at least one hour compared to wild-type HMCES (Figures 4D and 4E), suggesting that E127-dependent self-reversal is an important process in cells. After 2 hours of recovery, the DPC level was reduced almost to approximately the same amount in both cell lines, suggesting alternative mechanisms to complete removal in the absence of self-reversal.

**Figure 4.**
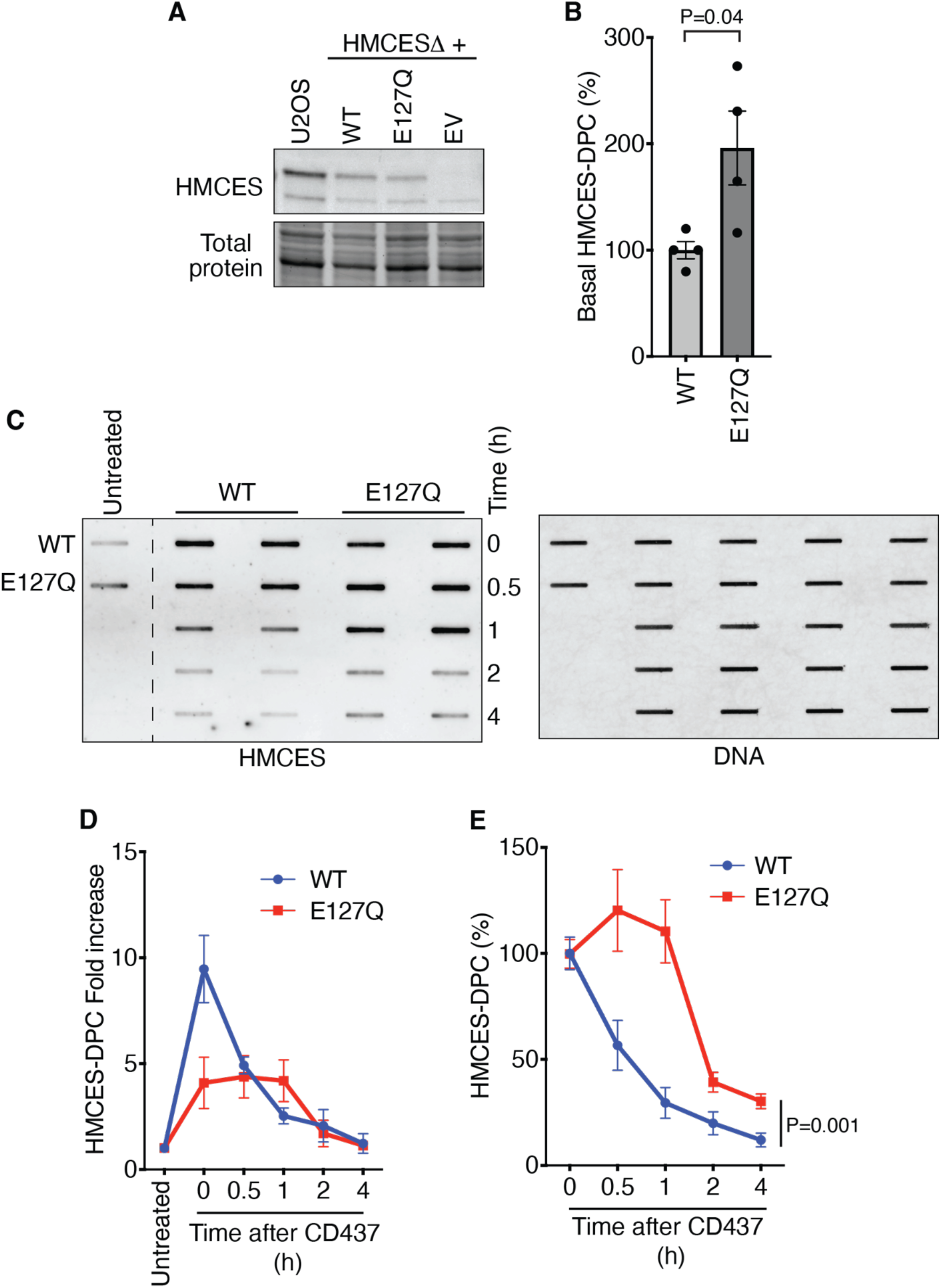
Inactivating HMCES self-reversal delays DPC resolution in cells. **(A)** Immunoblot of U2OS or HMCESΔ cells complemented with WT HMCES, E127Q HMCES, or empty vector (EV). **(B)** Quantification of HMCES-DPC levels in untreated cells that express WT or E127Q HMCES. Mean ± SEM, n=4, two-tailed t-test. **(C)** Representative image of RADAR assay of HMCES-DPC removal. Synchronized cells expressing WT or E127Q HMCES not exposed (untreated) or exposed to CD437 for 30 minutes followed by the indicated recovery times. **(D)** Quantification of HMCES-DPC removal. Analysis of fold-increase HMCES-DPC signal over time. Mean ± SEM, n=4 **(E)** Quantification of HMCES-DPC with time zero immediately after CD437 set at 100%. Mean ± SEM, n=4, two-way ANOVA.

DPC accumulation is detrimental to the cells (Kühbacher and Duxin 2020). Hence, we analyzed how the expression of the E127Q HMCES protein affects cell fitness. Without the addition of exogenous DNA damaging agents, E127Q HMCES showed around 30% less viability measured by an alamarBlue assay than HMCESΔ cells transduced with an empty vector (EV) or expressing WT HMCES (Figures 5A and S3). Moreover, over-expressing E127Q from a strong promoter (OE E127Q) had a more deleterious effect leading to substantially slower growth (Figures 5A-C and S3). In contrast, cells overexpressing WT HMCES (OE WT) or HMCESΔ cells had near-normal growth rates (Figures 5A-C and S3). This result suggests that an excess of HMCES that cannot undergo self-reversal after DPC formation reduces cell fitness even in the absence of added DNA damage.

**Figure 5.**
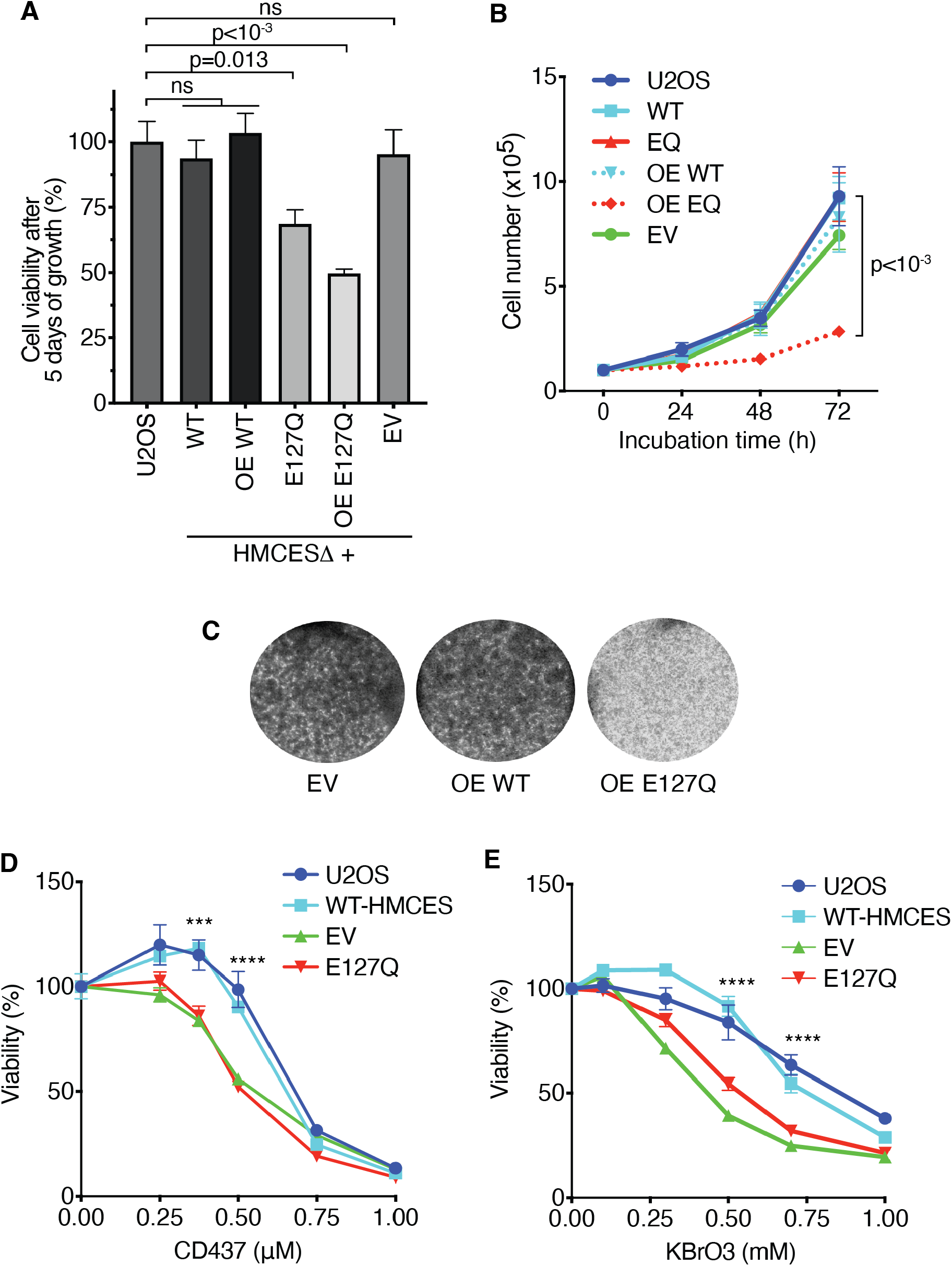
HMCES self-reversal is important for cell fitness and responses to DNA damage. **(A)** Cell viability measured using alamarBlue of the indicated cell lines measured 5 days after plating equal cell numbers. (OE = overexpression; EV = empty vector) Mean ± SEM, n=6, one-way ANOVA. **(B)** Cell proliferation analysis counting viable cells at each time point. Mean ± SEM, n=6, two-way ANOVA. **(C)** HMCESΔ cells overexpressing WT or E127Q HMCES or transduced with empty vector (EV) were seeded at equal cell numbers in 6-well dishes and incubated in normal growth media for 5 days before staining with methylene blue. **(D)** Percent viability of the indicated cells treated with CD437 as measured by metabolic activity four days after a 24-hour exposure to drug. Two-way ANOVA, n=3. WT vs E127Q, p<0.0004 (***), p<0.0001 (****). **(E)** Percent viability of the indicated cells treated with KBrO_3_ as measured by metabolic activity three days after a 48-hour exposure to drug. Two-way ANOVA, n=3. WT vs E127Q, p<0.0001 (****).

HMCESΔ cells and cells that express crosslink-deficient HMCES mutants are hypersensitive to DNA damage agents that generate AP sites (Mohni et al. 2019; Biayna et al. 2021; Mehta et al. 2020). To test whether inactivating self-reversal also causes hypersensitivity, we exposed cells expressing near endogenous levels of WT HMCES or the E127Q mutant to increasing doses of CD437 and let them recover in normal growth media before measuring viability. As expected, HMCESΔ cells transduced with an empty vector (EV) had reduced viability in the presence of CD437. E127Q cells also showed the same detrimental effect, meanwhile, WT HMCES recapitulated the viability of control U2OS cells (Figure 5D). Similarly, EV and E127Q cells were hypersensitive to KBrO_3_ (Figure 5E). These results further support the idea that HMCES self-reversal is an important mechanism for resolving the HMCES-DPC.

## DISCUSSION

Abasic sites are frequent DNA lesions that threaten genome stability especially when they are present in ssDNA where BER cannot be used for repair. HMCES provides an evolutionarily conserved mechanism to recognize and shield these ssDNA lesions from inappropriate processing that can generate DSBs. However, repair requires the removal of the HMCES-DPC. Our results demonstrate that there is a self-reversal mechanism in human cells. The reversibility of the crosslink depends on E127, which is positioned adjacent to the thiazolidine linkage created by the N-terminal cysteine residue (Thompson et al. 2019; Wang et al. 2019; Halabelian et al. 2019). Mutation of E127 largely prevents the reversal reaction without preventing crosslink formation in biochemical assays and delays the resolution of the HMCES-DPC in cells. Furthermore, cells expressing only the E127Q HMCES protein are hypersensitive to agents that increase AP site formation and overexpression of this mutant protein decreases cell viability.

We cannot rule out the possibility that some of the effects of E127Q HMCES in cells could be due to a reduction in the rate of crosslink formation. While the Glu127 residue is essential for the reversal reaction, in YedK the equivalent residue promotes ring opening of the AP site deoxyribose ring from the furan to aldehyde form to facilitate crosslink formation (Paulin, Cortez, and Eichman 2022). We did observe a reduced increase in the amount of E127Q HMCES-DPC in response to CD437 compared to wild-type HMCES. This difference is largely due to an increase in the basal level of the E127Q HMCES-DPC compared to wild-type in the untreated cells but could also reflect a reduced rate of crosslink formation. Nonetheless, the severe reduction in cell growth after E127Q HMCES overexpression is not observed in HMCESΔ cells or after overexpression of WT HMCES suggesting self-reversal is important.

Our data are consistent with recent biochemical studies showing that the thiazolidine linkage is reversible (Sugimoto et al. 2023; Donsbach et al. 2022; Paulin, Cortez, and Eichman 2022). This linkage was initially thought to be highly stable because it appeared unchanged in biochemical reactions even days after formation, could be observed by crystallography, and blocked the action of AP endonucleases (Thompson et al. 2019). However, the addition of a second ssDNA-AP oligonucleotide trap revealed that HMCES could move to another substrate (Paulin, Cortez, and Eichman 2022), suggesting that DNA binding surface and the catalytic pocket are unchanged after reversal.

Inhibiting HMCES-DPC self-reversal delayed but did not prevent DPC removal. Previous studies on DPC repair in *Xenopus* egg extracts found two major repair mechanisms involving the protease SPRTN and the proteasome (Larsen et al. 2019). HMCES is ubiquitylated (Mohni et al. 2019; Gallina et al. 2020); however, our results showed that blocking the proteasome or the E1 enzyme required for ubiquitylation had only small effects on the amount of HMCES-DPC removal at least in the CD437 system. SPRTN removes DPCs at replication forks (Vaz et al. 2016; Stingele et al. 2016), showing full activity when these bulky lesions are located at ss/dsDNA junctions (Reinking et al. 2020; Li et al. 2019). SPRTN can remove HMCES-DPCs formed as an intermediate in inter-strand crosslink repair (Semlow, MacKrell, and Walter 2022). However, in response to CD437, SPRTN inactivation did not stabilize the HMCES-DPC, possibly because an appropriate substrate is not generated even after CD437 removal. In addition to SPRTN and the proteasome, proteases including GCNA (Borgermann et al. 2019), FAM111A (Kojima et al. 2020), and DDI1 (Yip, Bodnar, and Rapoport 2020) can participate in DPC repair. Further studies will be needed to understand if any of these mechanisms contribute to the resolution of the HMCES-DPC in cells.

The DNA binding cleft in HMCES and YedK only accommodates ssDNA on one side of the AP site (Thompson et al. 2019; Wang et al. 2019; Halabelian et al. 2019; Amidon and Eichman 2020). This creates specificity for either AP sites in ssDNA or the ones that exist at a dsDNA-ssDNA junction such as what would form if a polymerase stalls at the lesion. The apparent rate of self-reversal is greatly increased when a complementary oligonucleotide is added to the HMCES-DPC to generate duplex DNA. This increased reversal rate is likely due to the inability of HMCES to rebind the duplex DNA that forms as HMCES releases from the ssDNA. Coupling DPC resolution to the generation of duplex DNA in cells would allow HMCES to shield the AP site until it can be properly repaired by BER (Figure 6). Duplex DNA formation in cells could be generated by TLS synthesis across from the HMCES-DPC. Recent studies showed that TLS across from the DPC could be facilitated either by HMCES proteolysis (Semlow, MacKrell, and Walter 2022; Sugimoto et al. 2023) or by the action of FANCJ on the intact DPC (Yaneva et al. 2023). In either case, the outcome would likely be increased rates of mutation. However, HMCES-deficient cells have increased mutation rates, increased recruitment of TLS polymerases to replication forks, and exhibit synthetic lethality with TLS polymerase inactivation (Mehta et al. 2020; Srivastava et al. 2020; Mohni et al. 2019). Alternatively, template switching could be utilized to generate the duplex DNA, providing an error-free repair mechanism (Figure 6). Further studies will be needed to determine which of these mechanisms is preferred.

**Figure 6.**
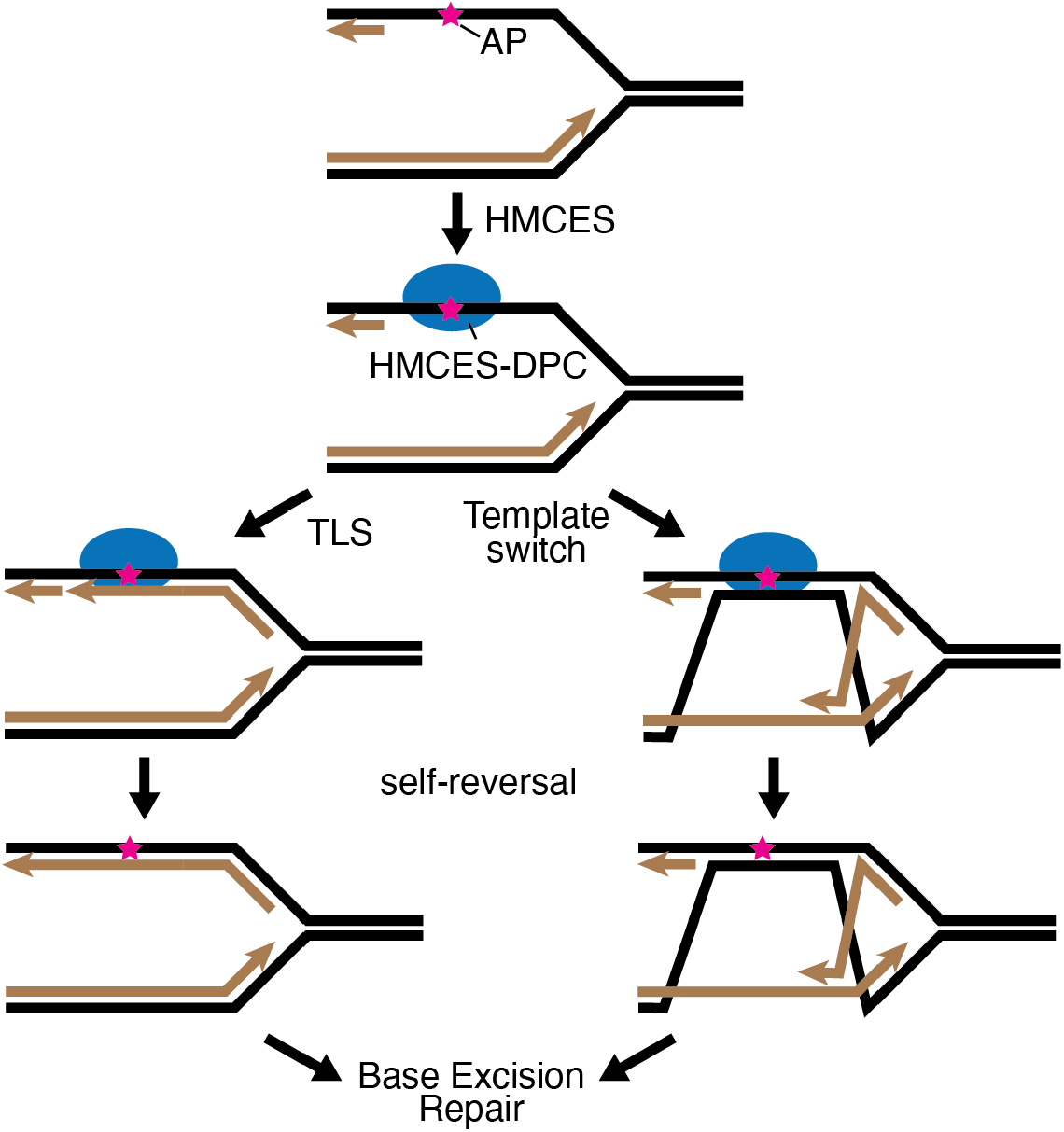
Model of CD437-induced HMCES-DPC reversal and repair.

Finally, to analyze HMCES-DPC resolution kinetics in cells we utilized POLα inhibition by CD437. Inhibiting POLα rapidly generates large amounts of ssDNA and HMCES-DPC formation. Although abasic sites form much more rapidly in ssDNA than dsDNA, further studies are needed to understand why CD437 is such an efficient generator of glycosylase-dependent HMCES-DPCs. In addition, the mechanisms that operate to remove the HMCES-DPC may differ depending on the context in which they are formed. Nonetheless, the synchrony in the formation and resolution of the HMCES-DPC after CD437 treatment provided a useful system to study how the HMCES-DPC is removed.

## ACKNOWLEDGEMENTS

We thank Dr. John Rouse for providing the SPRTN antibody. We thank Vanderbilt Antibody and Protein Resource for purification of the ssDNA antibody. The Vanderbilt Antibody and Protein Resource is supported by the Vanderbilt Institute of Chemical Biology and the Vanderbilt Ingram Cancer Center (P30CA68485). This work was supported by NIH grants R01ES030575 to D.C., R35GM136401 to B.F.E., K99ES034058 to K.P.M.M., and F31ES032334 to K.A.P.

## AUTHOR CONTRIBUTIONS

Conceptualization, J.R-F., B.F.E, and D.C.; Investigation, J.R-F., C.A.L., K.P.M.M., K.A.P, and Y.T; Writing-original draft, J.R-F. and D.C.; Writing-review and editing, J.R-F., C.A.L., K.P.M.M., K.A.P, Y.T., B.F.E., and D.C.; Funding Acquisition, D.C. and B.F.E.; Supervision, D.C. and B.F.E.

## AUTHOR INFORMATION

The authors have no conflicts of interest.

## METHODS

### Generation of stable cell lines

U2OS HMCESΔ cells were generated previously(Mohni et al. 2019). Stable cell lines expressing WT-HMCES, E127Q, or EV were generated by transduction of U2OS HMCESΔ cells with a pLPG backbone lentivirus. Stable cell lines overexpressing WT HMCES or E127Q were generated by transduction of U2OS HMCESΔ cells with pLPCX retrovirus containing a CMV promoter. Protein expression was corroborated by Western blot after three days of puromycin selection.

### Cell transfections

siRNA transfections were performed using Lipofectamine RNAimax according to manufacturer’s instructions.

### Plasmids

HMCES E127Q plasmid was generated by Gibson assembly of gene block containing the point mutation into the pLPG plasmid with gD promoter. Over-expression plasmids were created by cloning HMCES WT or E127Q cDNA into pLPCX plasmid containing CMV promoter. Plasmids were corroborated by sequencing. pUGI-NLS UDG Inhibitor (UGI) plasmid was purchased from Addgene (Cat#101091).

### Analysis of parental ssDNA with native BrdU staining

Cells were plated in a 96-well glass-bottom, poly-L-lysine coated plate and pulsed with 2mM BrdU for 18 hours. BrdU was washed off for two hours before drug treatment. Cells were treated with CD437 (5μM) or HU (0.3mM, 3mM) for 30 min and subsequently fixed with 3% paraformaldehyde, 2% sucrose for 10 min. Cells were permeabilized in PBT (0.5% triton X-100 in PBS) for 10 min, blocked for 1 hour in 10% normal goat serum, and probed with anti-BrdU antibody (AbCam Cat#ab6326), followed by Goat anti-Rat IgG (Thermo Cat# A11007). The plate was imaged and analyzed directly using a Molecular Devices ImageXpress high-content imager.

### AP site detection

Genomic DNA was purified as described in the RADAR method, quantified, and diluted to 100 ng/μl in dH_2_0. Abasic sites were labeled by incubation of 2.5 μg DNA with 5 mM biotinylated aldehyde reactive probe (ARP; Dojindo Laboratories, A305) for 1 hour at 37°C. The DNA was ethanol precipitated, washed twice with 70% ethanol, resuspended in dH_2_0, and quantified. For the loading control, 50 ng DNA was diluted in 6X SSC, denatured, and dot blotted onto a nylon membrane. The membrane was treated with 1.5 M NaCl, 0.5 N NaOH for 10 min, followed by 1 M NaCl, 0.5M Tris-HCl pH 7.0 for 10 min, and DNA was crosslinked to the membrane using a Stratalinker. The membrane was blocked with 5% milk in TBST immunoblotted for ssDNA (Millipore Sigma, MAB3034). For ARP detection, 500 ng or 250 ng DNA was diluted in 6X SSC, denatured, and applied to a nylon membrane, as above. The membrane was blocked with 5% bovine serum albumin in TBST and biotin was detected with Streptavidin-HRP (ThermoFisher).

### Immunoblotting

Cell lysates were extracted using Igepal lysis buffer (1% Igepal, 150mM NaCl, 50mM Tris pH 7.4) enriched with protease inhibitor cocktail (Roche). Proteins were analyzed by SDS-PAGE and immunoblotting.

### DNA combing

Cells were labeled for 20 min with 20 μM CldU (Sigma, C6891) followed by 40 min with 100 μM IdU (Sigma, l7125) and approximately 300,000 cells were collected by trypsinization and embedded in agarose plugs. DNA combing was performed according to the manufacturer’s instructions (Genomic Vision) with minor modifications using a combing machine. DNA-combed coverslips were baked for 2 hours at 65°C and stored at -20°C. The DNA-coated coverslips were denatured with freshly prepared 0.5M NaOH, 1M NaCl solution, washed with PBS, and dehydrated consecutively in 70%, 90%, and 100% ethanol before air drying. Coverslips were blocked with 10% goat serum, 0.1% Triton X-100 in 1X PBS and immunostaining was performed with antibodies that recognize CldU (Abcam, ab6326) and IdU (BD Biosciences, 347580) for 1 hour at room temperature. Coverslips were then washed in PBS, probed with secondary antibodies for 30 min at room temperature, washed with PBS and mounted using ProLong Gold (ThermoFisher). Images were captured using a 40X oil objective (Nikon Eclipse Ti) and fiber length analysis was performed using Nikon Elements software.

### RADAR assay

Cells were synchronized with 2mM thymidine overnight. Then, thymidine was removed and cells recovered in normal growth medium for 2h prior to treatment with 5 μM CD437 for 30 min. Cells were washed twice with PBS and recovered in normal growth medium. Cells were lysed in RADAR buffer (RLT plus buffer supplemented with 1% Sarkosyl). Genomic DNA was ethanol precipitated by the addition of ½ volume 100% ethanol and incubation at -20°C for 5 min. After full-speed centrifugation, the DNA pellet was washed twice with 70% ethanol and resuspended in 8mM NaOH at 65°C at 800rpm for 3 hr. DNA concentration was determined by spectrophotometry. DNA sample (20ug) was digested with Pierce universal nuclease in 1X TBS with 2mM MgCl_2_ at 37°C at 300rpm for 1 hr. Samples were boiled for 5 min and applied to a nitrocellulose membrane with a slot blot apparatus. The membrane was blocked for 1 hr with 5% non-fat dry milk in TBST and immunoblotted for HMCES. For the DNA blot, DNA sample (1 μg) was added in 1ml of 6X SSC buffer. The sample was boiled and then on ice for 10 min each and added to a nylon membrane with a slot apparatus. The membrane was placed face up on Whatman paper soaked with solution A (1.5M NaCl, 0.5M NaOH) for 10 min and in solution B (1.5M NaCl, 0.5M Tris pH7.5) for 5 min. After air drying, the membrane is crosslinked with UV 1200J/m^2^ and blocked for 1 hr with 5% non-fat dry milk in TBST and immunoblotted for ssDNA.

### Preparation of AP-DNA

Sequences of oligonucleotides used in the biochemical assays are listed in Supplementary Table 1. AP-DNA was prepared by incubation of 200 μM uracil-containing oligonucleotide (700 μM of the trap oligo) and 8 U UDG in UDG Buffer (supplemented with 1mM DTT) at 37°C for 20 min. AP-DNA was prepared fresh for each reaction.

### Cross-link reversal assay

WT and E127Q HMCES were purified as previously described(Paulin, Cortez, and Eichman 2022). HMCES-DPC was formed by incubation of 10 μM 20mer AP-DNA with 2 μM HMCES (WT or E127Q) in DPC buffer (20 μM HEPES, 10mM NaCl, 1mM EDT, pH 8) overnight at 37°C. DPC-20 was incubated with a 50-fold excess of 40mer AP-DNA to trap any reversed HMCES. Reactions were stopped by adding an equal volume of 2X SDS buffer. Each time point was initiated in reverse so that all reactions were quenched for the same length of time. Reaction products were resolved on 4 to 12% Bis-Tris gels and Coomassie stained for detection.

Reversal trapping in duplex DNA required incubation of 10 μM 40mer AP-DNA with 2 μM WT HMCES in DPC buffer overnight at 37°C. DPC-40 was incubated with a 1.4-fold excess of non-complementary oligo (40mer control) or the complementary oligo (c40mer). Simultaneously, a 50-fold excess of 20mer AP-DNA was added to trap any reversed HMCES. Each time point was initiated in reverse so all reactions were quenched for the same length of time and by adding an equal volume of 2X SDS buffer. Reactions products were resolved on 4 to 12% Bis-tris gels, 1X MES running buffer, and coomassie stained for protein. The reversal percentage was calculated as the percentage of DPC-20 compared to total at each time point.

### Viability assays

5×10^3^ cells/well in a 96-well dish were seeded for all the different cell lines. Cells were incubated in a DMEM growth medium for 5 days. For drug hypersensitivity assays, cells were treated with CD437 for 24 hr, or KBrO_3_ for 48 h and then recovered in normal growth medium for 4 and 3 days, respectively. AlamarBlue readout was performed using a BioTek multimode reader. All viability measurements are presented as a percentage of the untreated control. Overall cell growth of over-expression cell lines and EV was done by methylene blue staining. Cells were seeded and incubated for 5 days.

