## Supplemental figures for "Self-reversal facilitates the resolution of HMCES-DNA protein crosslinks in cells"

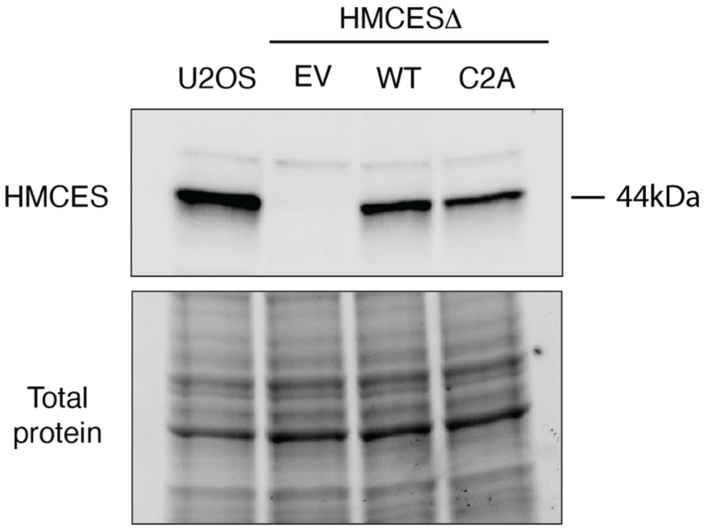


**Figures S1. Immunoblot of HMCES expression.**
Immunoblot analysis of total HMCES protein in U2OS cells and HMCESΔ cells complemented with empty vector (EV), WT, or C2A HMCES.

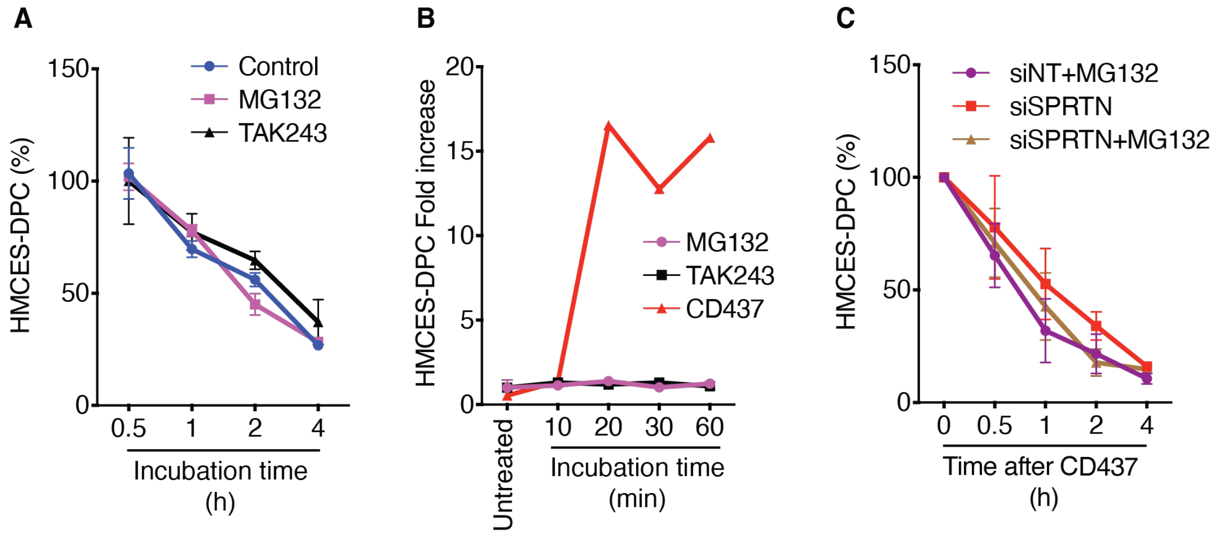


**Figures S2. Analysis of proteasome, E1 ubiquitin-activating enzyme, and SPRTN contributions to HMCES-DPC resolution.**

**(A)** Percentages remaining of HMCES-DPC with HMCES-DPC levels at 0.5h set to 100%. Mean ± SEM, n=3. **(B)** Quantification of HMCES-DPC formation during incubation with CD437, MG132, or TAK243 for the indicated times. **(C)** Quantification of HMCES-DPC removal in cells transfected with non-targeting (siNT) or SPRTN siRNAs and treated with MG132. MG132 was added during and after CD437 treatment.

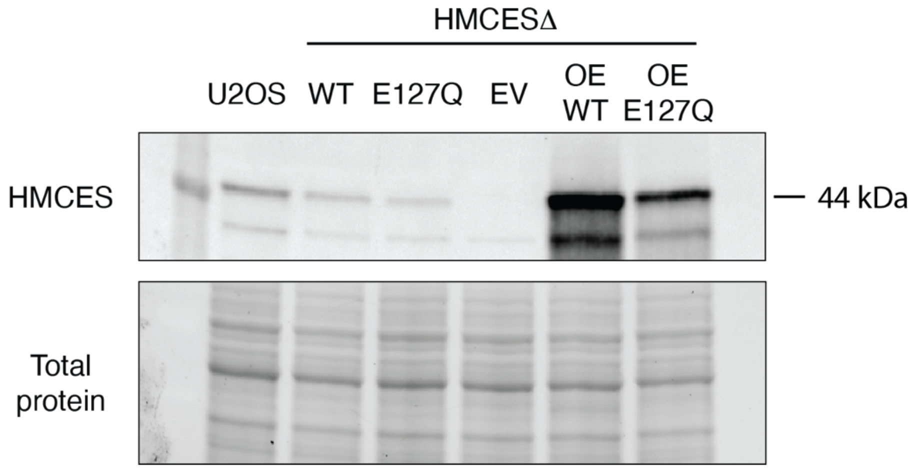


**Figures S3. Immunoblot of HMCES expression.**
Immunoblotting of U2OS or HMCESΔ cells complemented with WT, E127Q, or empty vector (EV) expressed from weak or strong promoters. OE = Overexpression.
